# Parent of Origin Effects on Quantitative Phenotypes in a Founder Population

**DOI:** 10.1101/150185

**Authors:** Sahar V. Mozaffari, Jeanne M. DeCara, Sanjiv J. Shah, Roberto M. Lang, Dan L. Nicolae, Carole Ober

## Abstract

The impact of the parental origin of associated alleles in GWAS has been largely ignored. Yet sequence variants could affect traits differently depending on whether they are inherited from the mother or the father. To explore this possibility, we studied 21 quantitative phenotypes in a large Hutterite pedigree. We first identified variants with significant single parent (maternal-only or paternal-only) effects, and then used a novel statistical model to identify variants with opposite parental effects. Overall, we identified parent of origin effects (POEs) on 11 phenotypes, most of which are risk factors for cardiovascular disease. Many of the loci with POEs have features of imprinted regions and many of the variants with POE are associated with the expression of nearby genes. Overall, our results indicate that POEs, which are often opposite in direction, are relatively common in humans, have potentially important clinical effects, and will be missed in traditional GWAS.

## INTRODUCTION

Genome-wide association studies (GWAS) typically treat alleles inherited from the mother and the father as equivalent, although variants can affect traits differently depending on whether they are maternal or paternal in origin. In particular, parent of origin effects (POEs) can result from imprinting, where epigenetic modifications allows for differential gene expression on homologous chromosomes that is determined by the parental origin of the chromosome. Mutations in imprinted genes or regions can result in diseases. For example, two very different diseases, Prader-Willi Syndrome (PWS) and Angelman Syndrome (AS), are due to loss of function alleles in genes within an imprinted region on chromosome 15q11-13. Inheriting a loss of function mutation for the *SNRPN* gene from the father results in PWS but inheriting a loss of function mutation for the *UBE3A* gene from the mother results in AS^1,2^. Long noncoding RNA genes at this and other imprinted regions act to silence (i.e. imprint) genes in *cis*. Imprinted genes are often part of imprinted gene networks, suggesting regulatory links between these genes^3^^-^^5^. More than 200 imprinted loci have been described in humans^6^, but there are likely many other, as yet undiscovered, imprinted loci.

Previous studies have utilized pedigrees to test maternal and paternal alleles separately for association with phenotypes or with gene expression to uncover new imprinted loci^6^^-^^9^. Kong *et al.* ^7^ discovered one locus associated with breast cancer risk only when the allele is inherited from the father and another locus associated with type 2 diabetes risk only when the allele is inherited from the mother. Garg *et al.* reported parent-of-origin *cis*-eQTLs with known or putative novel imprinted genes affecting gene expression^8^. Two additional studies by Zoledziewsk *et al.* and Benonisdottir *et al.* identified opposite POEs on adult height at known imprinted loci^6,10^. Both studies reported associations with variants at the *KCNQ1* gene, and one showed additional opposite POEs with height at two known imprinted loci (*IGF2-H19* and *DLK1-MEG3*)^6^. These studies provide proof-of-principle that alleles at imprinted loci can show POEs, some with opposite effects, with common phenotypes.

However, no previous study has included a broad range of human quantitative phenotypes or has considered whether genome-wide variants can have different effects depending on the parent of origin. To address this possibility, we developed a statistical model that directly compares the effects of the maternal and paternal alleles to identify effects that are different, including those that are opposite. We applied this model in a study of 21 common quantitative traits that were measured in the Hutterites, a founder population of European descent for which we have phased genotype data^11^. We identified variants with maternally inherited or paternally inherited effects only and variants with opposite POEs. Some of the identified regions have characteristics similar to known imprinted genes. Overall, we show that this model can identify putative novel imprinted regions with POEs for a broad range of clinically relevant quantitative phenotypes.

## RESULTS

### GWAS

We first performed standard genome-wide association studies (GWAS) of 21 traits in the Hutterites (**Supplementary Table 1**). These studies identified one genome wide significant association (p < 5×10^−8^) with each of five of the 21 traits: low density lipoprotein level (LDL)-cholesterol, triglycerides, carotid artery intima media thickness (CIMT), left ventricular mass index (LVMI), and monocyte count. The results of all 21 GWAS are summarized in **Supplementary Table 2** and **Supplementary Figs. 1** and **2**. Results for all variants for all GWAS are deposited in dbGaP (phs000185 – submission in progress).

### Parent of Origin GWAS

We considered two possible mechanisms of POEs. In the first, the effect size of one parent’s allele is close to zero and the effect size of the other parent’s allele is significantly different from zero. For these cases, we performed a paternal only or maternal only GWAS. In other cases, the maternal and paternal alleles may both have effect sizes different from zero, but the effects are significantly different from each other or opposite in direction. To detect these types of POEs, we developed a model that tests for differences between parental effects (see Methods). This model is especially powerful to identify variants with parental effects in opposite directions.

#### Maternal and Paternal GWAS

Using the same phenotypes, genotypes, pedigree, and criteria for significance as in the standard GWAS, we tested for maternal and paternal effects on each trait by testing each parentally inherited allele with the trait of interest, similar to previous studies^7,8,10^. Variants were considered to have POEs if they had a p-value less than 5×10^−8^ in only one parent and were not significant in the standard GWAS (i.e., the LDL association on chromosome 19 and the triglycerides association chromosome 11 were not considered to have POEs; see **Supplementary Table 1**). The most significant parent of origin associations are summarized in **Table 1**. All significant results of the parent of origin GWAS for all 21 phenotypes are included in **Supplementary Table 5**.

**Table 1.**
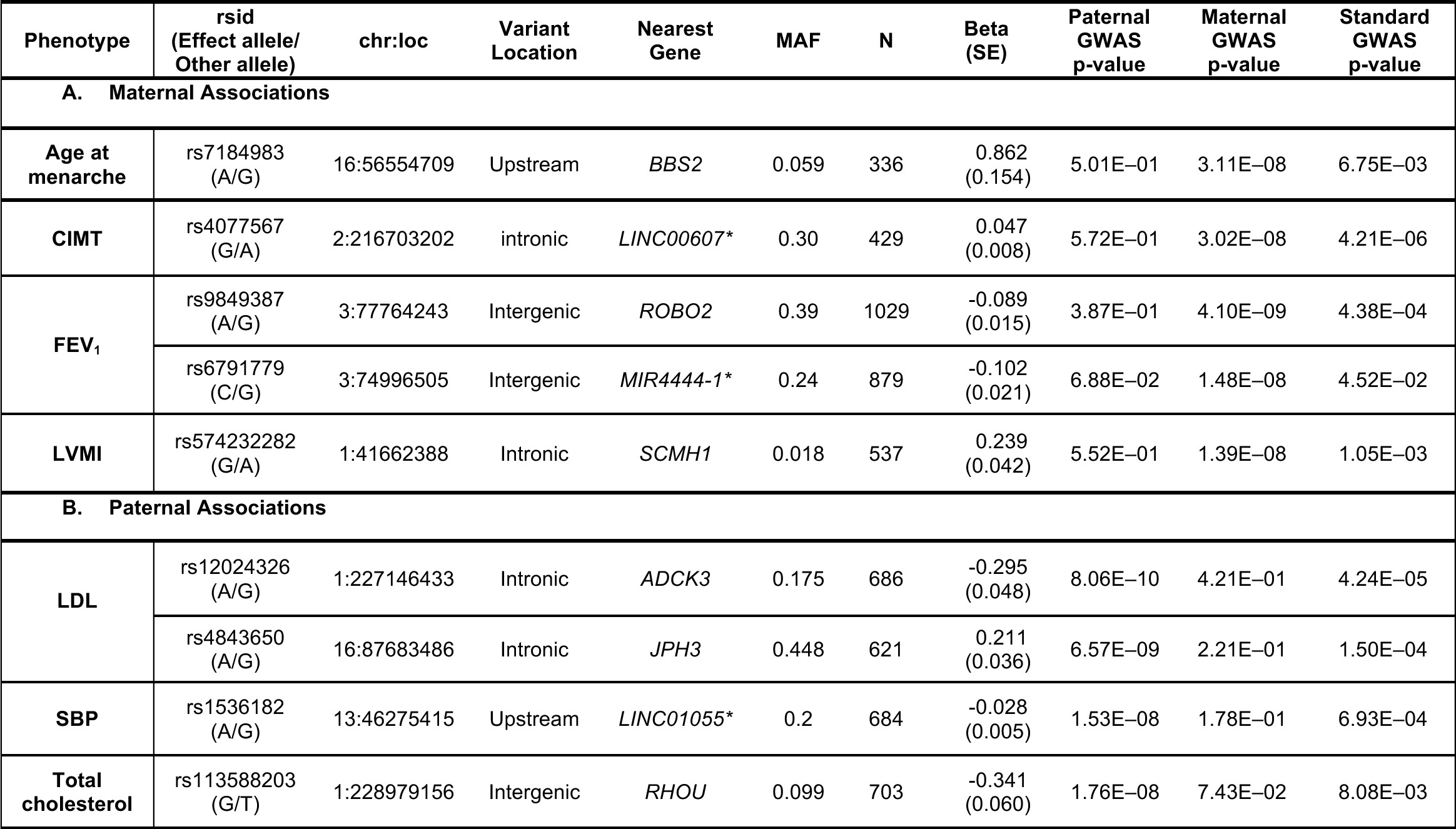
Phenotypes with significant single parent of origin associations. The most significant variant (P < 5×10^−8^) at each locus for the (A) maternal and (B) paternal associations associated with each phenotype is shown. *non-coding RNA genes

Overall, seven phenotypes had genome-wide significant parent of origin associations: four in the maternal only GWAS and three in the paternal only GWAS. Three cardiovascular disease (CVD)-associated phenotypes (age at menarche, CIMT, LVMI) and one lung function phenotype (forced expiratory volume in one second [FEV_1_]) were associated with maternally-inherited alleles only.

A maternally inherited allele at rs7184983 (G) on chromosome 16 was associated with younger age of menarche (P = 3.11 ×10^−8^) (**Fig. 1**). This SNP, rs7184983, is located upstream of the *BBS2* gene and is associated with increased expression of *OGFOD1* in transformed fibroblast cells and tibial nerve^12^. The maternally inherited allele at rs4077567 (G) on chromosome 2 was associated with decreased CIMT (P = 3.02×10^−8^) (**Supplementary Fig. 2**). This SNP is in the intron of a long intergenic noncoding gene, *LINC00607*, that is expressed in aorta, coronary, and tibial artery, all tissues potentially relevant to CIMT and atherosclerosis^12^. A maternally inherited allele at rs574232282 (G) in the intron of *SCMH1* on chromosome 1 was associated with increased LVMI (P = 1.39 ×10^−8^) (**Supplementary Fig. 3**). *SCMH1* is expressed in aorta, coronary, and tibial artery^12^. SCMH1 protein associates with the polycomb group multiprotein complexes required to maintain the transcriptionally repressive state of certain genes^12^. Lastly, maternally inherited alleles at rs9849387 (A) and rs6791779 (C) on chromosome 3 were both associated with reduced FEV_1_ (P= 4.10×10^−9^ and 1.48×10^−8^, respectively) (**Supplementary Fig. 4**). The nearest gene to rs9849387 is *ROBO2* (65kb, downstream), which is expressed in the lung as well as in brain, and ovary^12^. The nearest gene to rs6791779 is MIR4444-1(267kb) whose expression has not been characterized.

**Figure 1.**
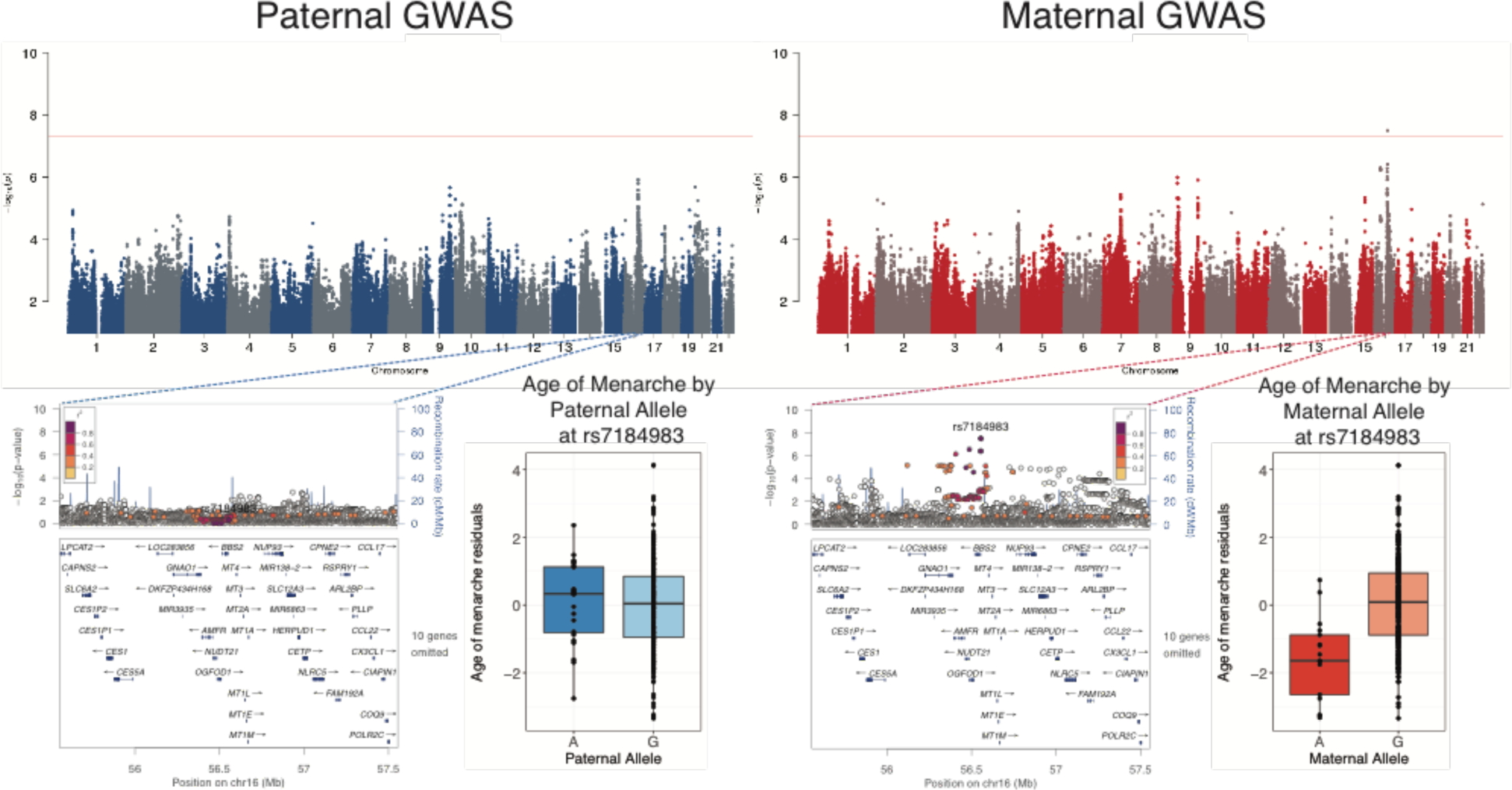
Maternal and Paternal GWAS results for Age of Menarche. The top panel shows the Manhattan plots from the maternal (left) and maternal (right) GWAS. LocusZoom plots for both GWAS are shown in the lower panel for the associated region in the GWAS. Boxplots show the distribution of age of menarche residuals (y-axes) by the corresponding maternal and paternal alleles at this SNP (x-axes). The horizontal bar of the boxplot shows the median, the box delineates the first and third quartile, and the whiskers show +/-1.5 x IQR.

**Figure 2.**
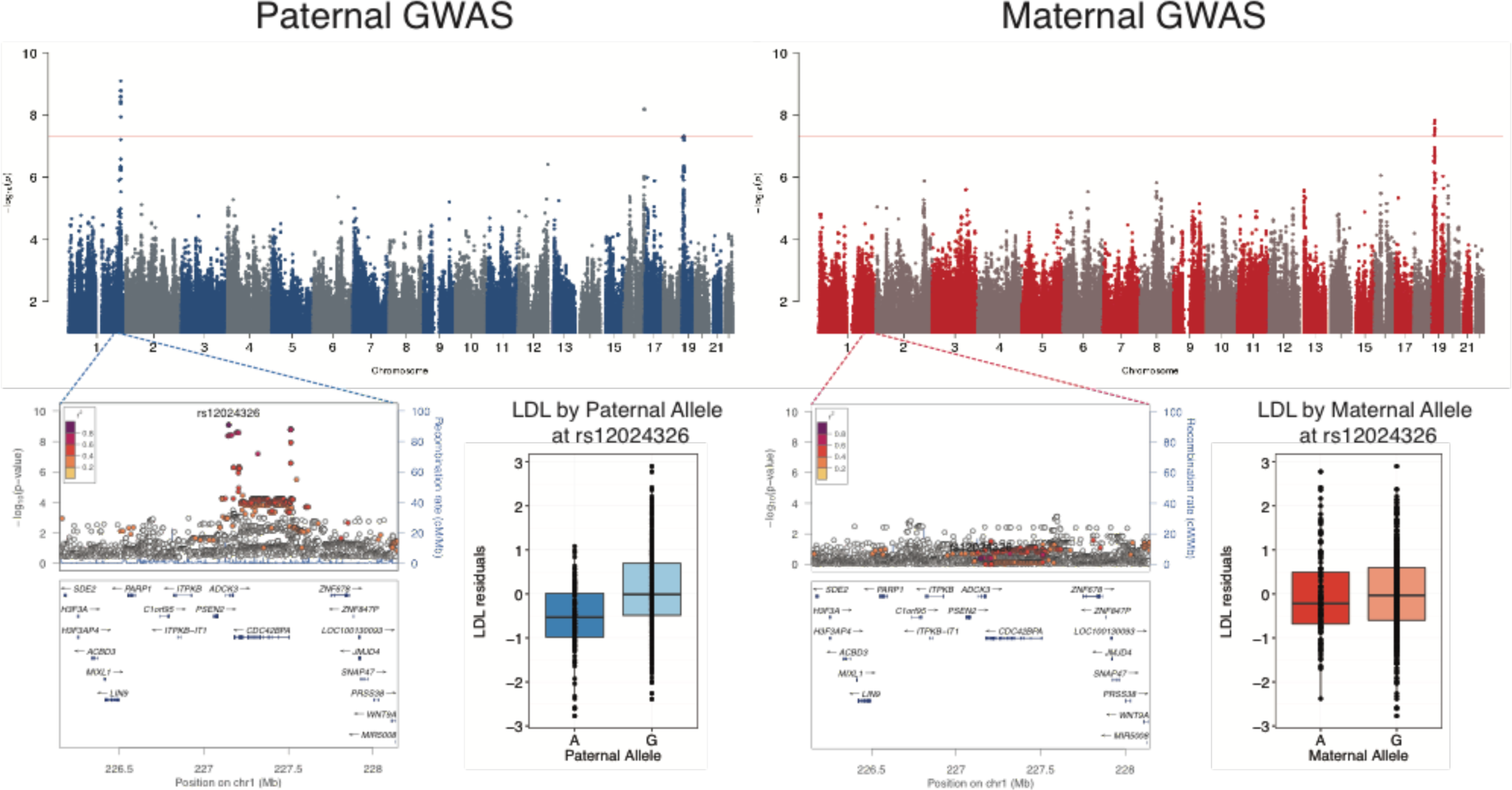
Maternal and Paternal GWAS results for LDL Cholesterol. The top panel shows the Manhattan plots from the maternal (left) and maternal (right) GWAS. LocusZoom plots for both GWAS are shown in the lower panel for the associated region in the GWAS. Boxplots show the distribution of LDL residuals (y-axes) by the corresponding maternal and paternal alleles at this SNP (x-axes). The horizontal bar of the boxplot shows the median, the box delineates the first and third quartile, and the whiskers show +/-1.5 x IQR.

Three other CVD-related phenotypes (systolic blood pressure, LDL-C, and total cholesterol) had associations with paternally inherited alleles only. The paternally inherited allele at rs12024326 (A) on chromosome 1 was associated with lower LDL-cholesterol levels (P = 8.06×10^−10^) (**Figure 2**). rs12024326 is in the intron of gene *ADCK3*, and the same allele was associated with increased expression of *ADCK3* in whole blood, as well as decreased expression of a neighboring gene, *CDC42BPA* in brain (cerebellum), heart (left ventricle), esophagus, and tibial artery ^12^. The paternal G allele at rs4843650 on chromosome 16 was associated with increased LDL-C and is located in the intron of *JPH3*, which is expressed predominantly in the brain^12^. A SNP on chromosome 13 (rs1536182) was associated with systolic blood pressure levels when it was inherited from the father (**Supplementary Fig. 5**). The paternally inherited A allele at this SNP was associated with decreased systolic blood pressure, as well as decreased expression of its closest gene, *LINC01055*, a long intergenic noncoding gene, in testis^12^. A paternally inherited allele at rs113588203 (G) on chromosome 1 was associated with lower total cholesterol (P = 1.76×10^−8^) (**Supplementary Fig. 6**). This SNP is intergenic between *RHOU* (96kb, downstream), which is expressed across multiple tissues, and *MIRR4454* (331kb), which is expressed in adipose, kidney and heart tissues^12^.

#### GWAS for Differential Parent of Origin Effects

Because some imprinted regions include genes that have both maternal or paternal specific tissue expression, we next tested for such differential effects with these 21 phenotypes. In these analyses, we compared the effect and direction of the association between maternal and paternal alleles to identify variants that have different effects, including opposite effects, on the phenotype. Such loci would be completely hidden in standard GWAS in which paternally and maternally inherited alleles are combined. These opposite effect GWAS revealed 11 independent loci with opposite POEs for nine different traits, at least six of which are associated with CVD risk (**Table 3**, **Supplementary Fig. 7**).

A locus on chromosome 16, near the *CDH8* gene (128kb, upstream), was associated with opposite POEs with age of menarche (**Fig. 3**). *CDH8* is highly expressed in the brain, as well as in the aorta artery and pituitary gland. Two loci on chromosomes 5 and 6 were associated with opposite POEs on body mass index (BMI) (**Fig. 4**). The most significant variant on chromosome 5 (rs77785972) is near a long intergenic noncoding gene, *LINC01340* (409kb, downstream), whose expression has not been well characterized. The SNP on chromosome 6 (rs17605739) is also in a long intergenic noncoding gene, *RP1-209A6.1*, which is expressed in low levels in the tibial artery, bladder, spleen, lung, pituitary gland, as well as testis.

**Figure 3.**
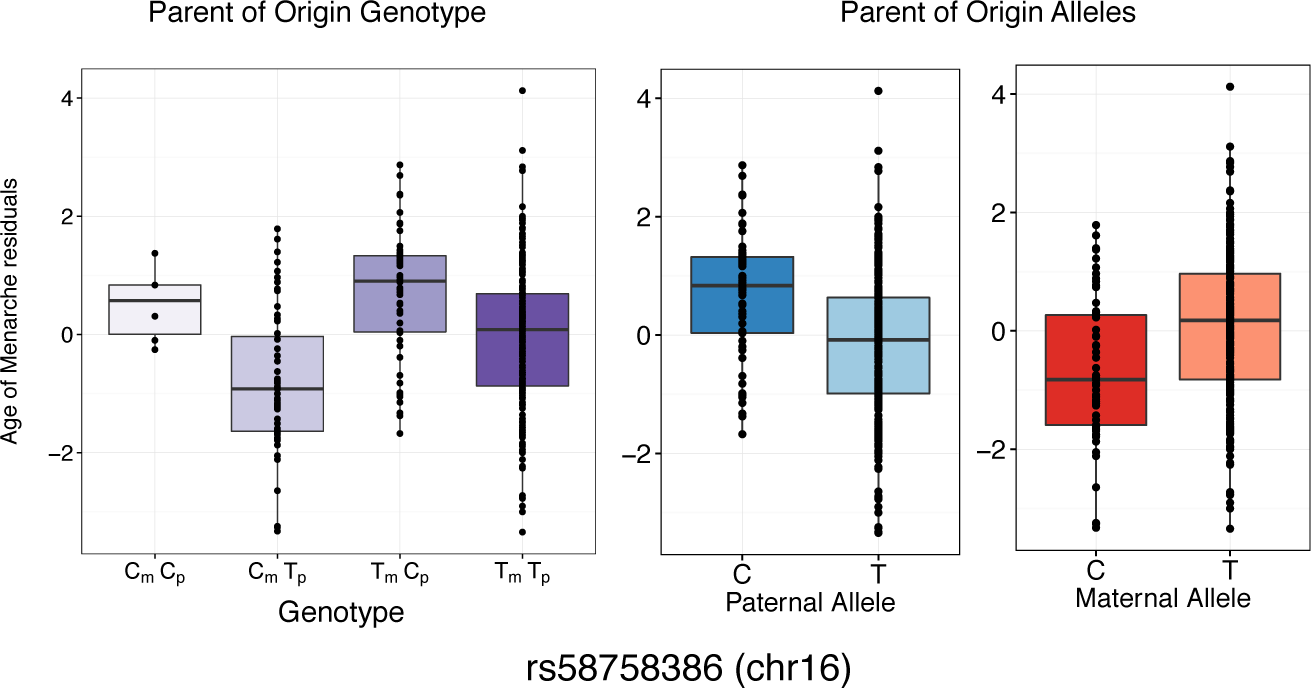
Opposite Effect Parent of Origin GWAS Result for Age of Menarche. Box plots of age of menarche residuals (y-axes) are shown for each of the four genotypes (left panel; x-axis), and for paternal (center panel; x-axis) and maternal (right panel; x-axis) alleles. The maternal C allele is associated with decreased and maternal T allele with increased age of menarche. The paternal C allele is associated with increased and the paternal T allele with decreased age of menarche. The horizontal bar of the boxplot shows the median, the box delineates the first and third quartile, and the whiskers show +/-1.5 x IQR.

**Figure 4.**
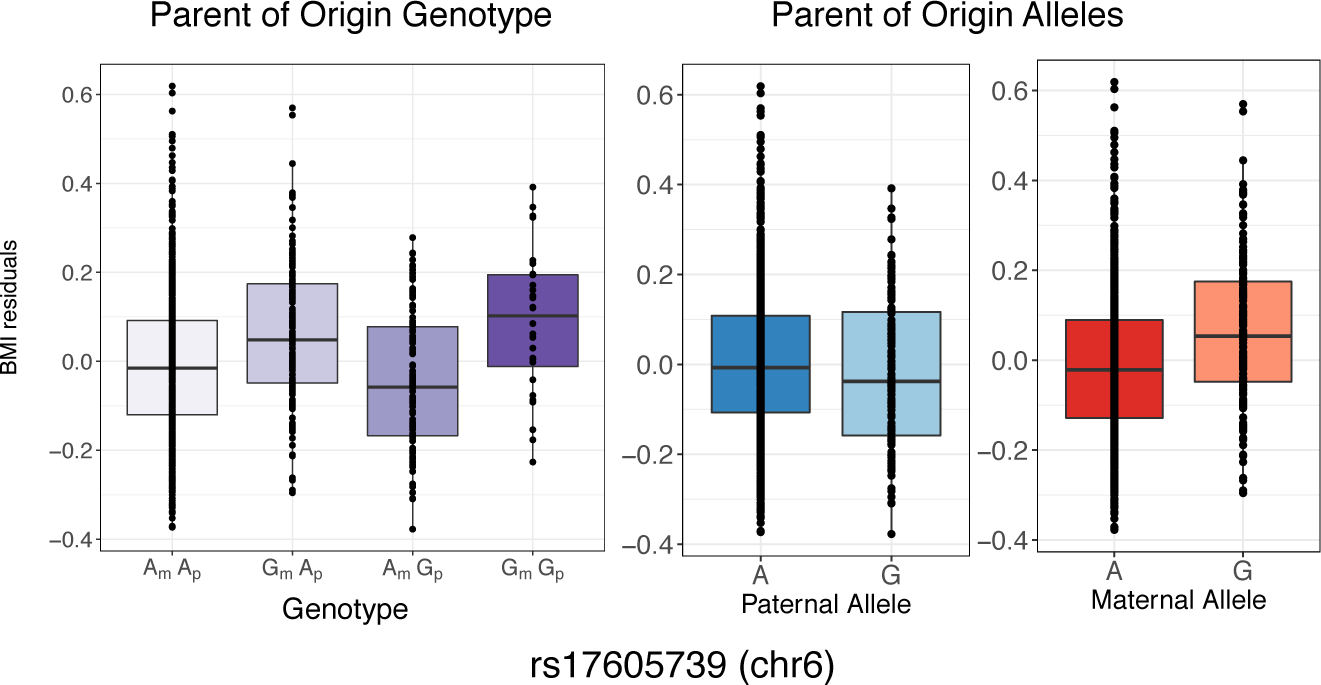
Opposite Effect Parent of Origin GWAS Result for BMI. Box plots of two significant loci plot BMI residuals (y-axes) for each of the four genotypes (left panel; x-axis), and for paternal (center panel; x-axis) and maternal (right panel; x-axis) alleles. For the **a.** SNP on chromosome 5 the maternal A allele is associated with decreased and maternal G allele with increased BMI. The paternal A allele is associated with increased and the paternal G allele with decreased BMI. For the **b.** SNP on chromosome 6 the maternal A allele is associated with decreased and maternal G allele with increased BMI. The paternal A allele is associated with increased and the paternal G allele with decreased BMI. The horizontal bar of the boxplot shows the median, the box delineates the first and third quartile, and the whiskers show +/-1.5 x IQR.

A SNP on chromosome 16 (rs1032596) was associated with opposite POEs on LDL-cholesterol (**Supplementary Fig. 8**). This SNP lies in the intron of another long noncoding RNA gene, *LINC01081*, which has been suggested to be imprinted because its downstream genes have also been shown to have parent- and tissue-specific activity^13^. A region on chromosome 2 has opposite effects associated with LVMI (**Supplementary Fig. 9**). The associated SNPs are in the intron of *XIRP2*, a cardiomyopathy associated protein that is expressed in skeletal muscle and heart left ventricle, suggesting that this gene could play a role in determining left ventricular mass^12,14,15^. In addition, the most significant SNP at this region, rs17616252 (and multiple SNPs in LD) is a strong eQTL (P = 1.8×10^−13^) for the gene *XIRP2* in skeletal muscle, *XIRP2-AS1* in testis, and *B3GALT1* in transformed fibroblast cells^12^. Four variants in a region on chromosome 1 in a microRNA gene, *MIR548F3*, were associated with opposite POEs on triglyceride levels (**Supplementary Fig. 10**). The expression of *MIR548F3* has not been characterized. SNP rs7033776 near *MELK* (27kb, downstream) on chromosome 9 was associated with opposite effects on total cholesterol (**Supplementary Fig. 11**). *MELK* is expressed in the colon and esophagus in addition to transformed lymphocytes and fibroblasts^12^.

Nine linked variants on chromosome 1 were associated with opposite POEs of blood eosinophil count (**Supplementary Fig. 12**). These variants are near the gene *IGSF21* (27kb, downstream) which is a member of the immunoglobulin superfamily and likely acts as a receptor in immune response pathways^16^. A variant on chromosome 3, rs12714812, was associated with opposite POEs for FEV_1_ (**Supplementary Fig. 13**). This variant has been shown to regulate the expression of a gene *CNTN3* (45kb, upstream) in heart and brain^12^. Studies in mice have suggested that this gene is imprinted and maternally expressed in the murine placenta^17^. Variant rs142030841 in the intron of the gene *TPGS2* on chromosome 18 has opposite POEs with neutrophil levels (**Supplementary Fig. 14**). This SNP is an expression quantitative trait locus (eQTL) for the noncoding RNA gene *RP11-95O2.5* in skin, testis, breast, thyroid and adipose tissue, for *CELF4* in tibial nerve and lung, and for *TPGS2* in tibial artery and transformed fibroblast cells^12^.

### Parent of Origin Effects on Gene Expression

To determine if any of the associated variants also showed POEs on gene expression in the Hutterites, we used RNA-seq gene expression data from lymphoblastoid cell lines (LCLs) collected from 430 of the individuals in the GWAS sample. We first tested for association of maternal and paternal variants with genes detected as expressed in the LCLs and whose transcript start site was within 1Mb of each associated SNP. There were no significant associations after multiple testing correction, similar to a previous study^6^. However, because we considered this to be exploratory analyses, we show results for the five most significant parent of origin eQTLs (**Table 3**). We next used the opposite effect model for each SNP in Table 2 and expression of all genes that were detected as expressed in LCLs and whose transcript start site was within 1Mb of the associated SNP. This resulted in 57 tests (1 SNP for each of 8 phenotypes, and 57 genes). The five most significant opposite effect eQTLs, none of which passed the Bonferroni threshold of 8.77 x 10^−4^, are shown in **Table 4**. The most significant opposite effect eQTL was for *POLR1E* expression with the SNP on chromosome 9 (rs7033776) that was associated with total cholesterol (opposite effect eQTL P = 9.86 x 10^−4^) (**Supplementary Fig. 15**). *POLR1E* is involved in the purine metabolism pathway as well as DNA-directed polymerase activity. The same SNP, rs7033776, had modest opposite effects with the expression of three other genes in the region (*PAX5*, *FBXO10*, and *FRMPD1*), a signature consistent with an imprinted region. Another SNP with opposite POEs on LVMI, rs16853098, was an opposite effect eQTL for *STK39*, a gene that has been previously associated with hypertension^18^.

**Table 2.** Significant Opposite Parent of Origin Effect GWAS Associations. The most significant variant at each locus for each phenotype is shown. β_M_ – β_P_ represents difference in parental effect size. *non-coding RNA genes

| Phenotype | rsid | chr:loc | Variant Location | Nearest Gene | MAF | $\beta_M - \beta_P$ (SE) | Opposite Effect GWAS | Paternal GWAS | | Maternal GWAS | | Standard GWAS p-value |
| --- | --- | --- | --- | --- | --- | --- | --- | --- | --- | --- | --- | --- |
|  |  |  |  |  |  |  |  | P-value | Beta (SE) | P-value | Beta (SE) |  |
| Age of menarche | rs12447191 | 16:62199299 | intergenic | *CDH8* | 0.17 | -0.654 (0.109) | 5.27E-09 | 5.20E-06 | 0.391 (0.085) | 1.85E-05 | -0.368 (0.085) | 8.68E-01 |
| BMI | rs77785972 | 5:97415767 | intergenic | *LINC01340\** | 0.025 | 0.154 (0.025) | 5.12E-10 | 5.84E-07 | -0.094 (0.019) | 1.58E-05 | 0.081 (0.019) | 5.39E-01 |
|  | rs17605739 | 6:22962798 | intronic | *RP1-209A6.1\** | 0.17 | 0.053 (0.010) | 3.01E-08 | 6.99E-05 | -0.032 (0.008) | 1.42E-06 | 0.034 (0.007) | 1.56E-01 |
| eosinophil | rs2355879 | 1:18732860 | intergenic | *IGSF21* | 0.14 | 0.091 (0.016) | 1.69E-08 | 5.83E-08 | -0.065 (0.012) | 5.59E-04 | 0.043 (0.012) | 2.53E-01 |
| FEV<sub>1</sub> | rs12714812 | 3:74813002 | intergenic | *CNTN3* | 0.45 | -0.119 (0.021) | 4.52E-08 | 1.78E-03 | 0.052 (0.017) | 6.35E-06 | -0.073 (0.016) | 9.58E-01 |
| LDL | rs1032596 | 16:86281537 | Intronic | *LINC01081\** | 0.30 | -0.310 (0.056) | 3.69E-08 | 1.05E-06 | 0.201 (0.041) | 4.56E-04 | -0.148 (0.042) | 2.71E-01 |
| LVMI | rs16853098 | 2:168013281 | intronic | *XIRP2* | 0.12 | -0.091 (0.053) | 4.18E-08 | 5.29E-06 | 0.064 (0.014) | 2.04E-04 | -0.048 (0.013) | 9.26E-01 |
| neutrophils | rs142030841 | 18:34371947 | intronic | *TPGS2* | 0.042 | -0.224 (0.041) | 4.40E-08 | 2.25E-03 | 0.078 (0.025) | 1.30E-07 | -0.188 (0.035) | 5.77E-01 |
| Triglycerides | rs7525463 | 1:218860879 | intronic | *MIR548F3\** | 0.16 | -0.401 (0.071) | 2.51E-08 | 1.14E-03 | 0.195 (0.060) | 5.52E-08 | -0.267 (0.049) | 2.84E-02 |
| Total cholesterol | rs7033776 | 9:36704465 | intergenic | *MELK* | 0.41 | 0.230 (0.041) | 4.12E-08 | 5.60E-08 | -0.183 (0.034) | 2.28E-03 | 0.099 (0.032) | 6.70E-02 |

**Table 3.**
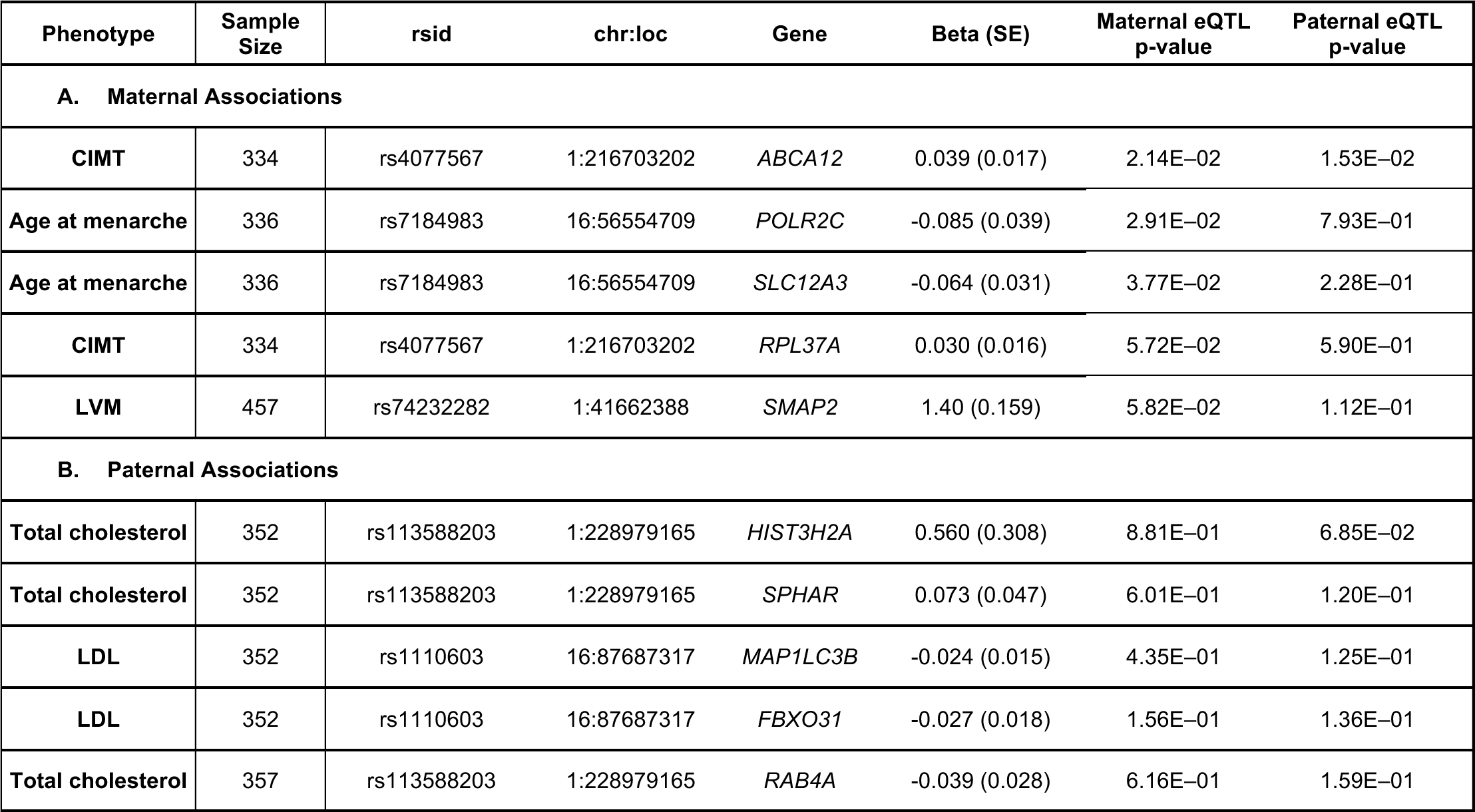
Parent of Origin eQTLs in LCLs. The most significant SNP for each phenotype (**Table 1**) was tested for association with gene expression for genes with TSS within 1Mb of the SNP. The effect sizes correspond to the maternal (A) or paternal (B) effect sizes.

| Phenotype | Sample Size | rsid | chr:loc | Gene | Beta (SE) | Maternal eQTL p-value | Paternal eQTL p-value |
| --- | --- | --- | --- | --- | --- | --- | --- |
| **A. Maternal Associations** |  |  |  |  |  |  |  |
| **CIMT** | 334 | rs4077567 | 1:216703202 | *ABCA12* | 0.039 (0.017) | 2.14E-02 | 1.53E-02 |
| **Age at menarche** | 336 | rs7184983 | 16:56554709 | *POLR2C* | -0.085 (0.039) | 2.91E-02 | 7.93E-01 |
| **Age at menarche** | 336 | rs7184983 | 16:56554709 | *SLC12A3* | -0.064 (0.031) | 3.77E-02 | 2.28E-01 |
| **CIMT** | 334 | rs4077567 | 1:216703202 | *RPL37A* | 0.030 (0.016) | 5.72E-02 | 5.90E-01 |
| **LVM** | 457 | rs74232282 | 1:41662388 | *SMAP2* | 1.40 (0.159) | 5.82E-02 | 1.12E-01 |
| **B. Paternal Associations** |  |  |  |  |  |  |  |
| **Total cholesterol** | 352 | rs113588203 | 1:228979165 | *HIST3H2A* | 0.560 (0.308) | 8.81E-01 | 6.85E-02 |
| **Total cholesterol** | 352 | rs113588203 | 1:228979165 | *SPHAR* | 0.073 (0.047) | 6.01E-01 | 1.20E-01 |
| **LDL** | 352 | rs1110603 | 16:87687317 | *MAP1LC3B* | -0.024 (0.015) | 4.35E-01 | 1.25E-01 |
| **LDL** | 352 | rs1110603 | 16:87687317 | *FBXO31* | -0.027 (0.018) | 1.56E-01 | 1.36E-01 |
| **Total cholesterol** | 357 | rs113588203 | 1:228979165 | *RAB4A* | -0.039 (0.028) | 6.16E-01 | 1.59E-01 |

**Table 4.**
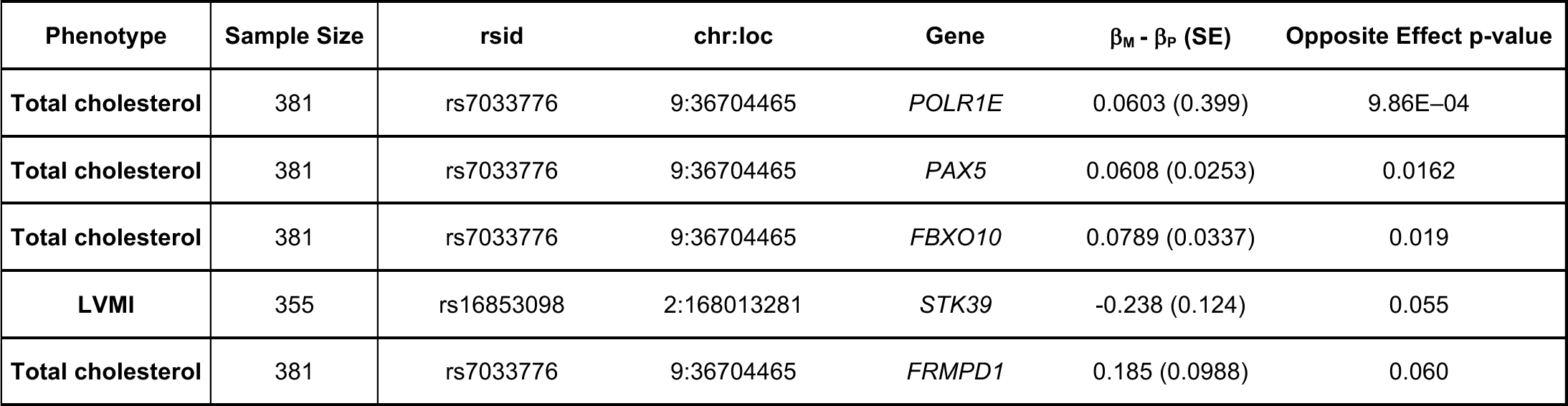
Opposite Parent of Origin eQTLs in LCLs. The most significant SNP for each phenotype (**Table 2**) was tested for opposite effect eQTLs with genes with TSS within 1Mb of the SNP. The effect size corresponds to the difference in maternal and paternal effect sizes.

## DISCUSSION

In this study, we introduced a novel statistical method that allows assessment of standard GWAS signals along with measures of differential POEs on common quantitative phenotypes. Similar to previous parent of origin studies of fewer phenotypes, we tested for associations of maternally- or paternally-derived alleles with each phenotype. We then extended this method to identify variants for which maternally- and paternally-derived alleles have different, including opposite, effects on phenotypic values. The focus on 21 common disease-associated phenotypes in a single large pedigree allowed us to broadly survey physiological effects of putative imprinted regions and the candidate genes at each associated locus. In contrast to previous studies, our new model can identify variants with opposite POEs that would be missed in traditional GWAS (**Table 2**).

Our studies of >1,000 Hutterites who are related to each other in a single pedigree allowed us to detect POEs, even when few genome-wide significant associations were detected in standard GWAS of the same phenotypes. Our method revealed parent of origin specific genome-wide significant associations for seven of the 21 phenotypes examined, with maternally-inherited alleles associated with four phenotypes, paternally-inherited alleles with three phenotypes (**Table 1**), and opposite parent of origin alleles with nine phenotypes, of which five also showed single POEs at different loci (**Table 2**). Overall, 11 of the 21 phenotypes examined showed genome-wide significant evidence of POEs with alleles at one or more loci. In contrast, standard GWAS of these same phenotypes and using the same markers in these same individuals revealed genome-wide significant association for only five traits.

It is notable that four of the nine significant opposite parent of origin effects (one each with LDL-C and triglycerides, and two with BMI) lie in or near long intergenic non-coding RNA genes (lincRNAs). LincRNAs are a feature of imprinted regions^1^, where they can silence the expression of genes on the opposite chromosome^3,19^. One of the variants, rs1032596, with an opposite parent of origin effect on LDL-C is located in the *LINC01081* gene. This noncoding RNA, along with *LINC01082*, regulates the *FOXF1* enhancer resulting in *FOXF1* parent- and tissue-specific activity^13^ providing experimental support for tissue specific expression, a feature of imprinted regions.

Another variant with POEs in our study has been suggested to be imprinted in previously published work. The variant associated with opposite POEs for FEV_1_ is an eQTL for the gene *CNTN3*. *CNTN3* was shown to have exclusive maternal allele-specific expression in murine placentas^17^, although this finding may have been due to contaminating maternal cells^20,21^.

Other regions associated with POEs harbor genes involved in transcriptional repression (e.g., *SCMH1* with LVMI on chromosome 1) or the associated SNPs are reported as eQTLs in GTEx with expression in tissues relevant to the phenotype under investigation (e.g., the LVMI-associated SNPs are eQTLs for *XIRP2*, which is expressed in skeletal muscle and heart left ventricle)^12^. Overall, these patterns of expression provide additional support that the parent of origin associations in our study are flagging imprinted regions or regions involved in the regulation of gene expression. Finally, we used gene expression in LCLs from the Hutterites to directly test for parent of origin eQTLs among SNPs associated with phenotypes in the parent of origin GWAS. Although none of the parent of origin eQTLs met criteria for significance after correcting for multiple testing, the SNP on chromosome 9 with opposite POEs on total cholesterol levels was borderline significant as an opposite parent of origin eQTL for *POLR1E*, and possible for three other genes at the same locus (*PAX5*, *FBXO10*, and *FRMPD1*). The presence of multiple genes with potential parent of origin expression patterns is further supportive of an imprinted locus. The availability of gene expression only in LCLs from the Hutterites limits the inferences we can draw about effects on expression because imprinted regions are often tissue-specific and sometimes developmentally regulated^1,2^. Despite this limitation, the fact that many of the SNPs associated with POEs on phenotypes are themselves eQTLs in relevant tissues (GTEx) and some are suggestive of having POEs on expression in LCLs from the Hutterites is generally supportive of the suggestion that some of the regions identified in this study are imprinted in humans. Additionally, our data suggest that loci with POEs influence a broad spectrum of quantitative phenotypes that are themselves risk factors for common diseases.

In particular, the discovery of POEs for eight traits that are associated cardiovascular disease risk is intriguing. These include metabolic phenotypes, such as BMI, total cholesterol, triglycerides, LDL, and age of menarche, that have indirect effects on cardiac health, as well as LVMI and CIMT, which more directly reflect cardiac health. Some of these phenotypes showed associations with paternally inherited alleles only (systolic blood pressure, LDL-C, total cholesterol), maternally inherited alleles only (LVMI, CIMT, and age at menarche), and/or with opposite effect variants (BMI, LDL-C, triglycerides, total cholesterol, LVMI, age at menarche). It has been suggested that genomic imprinting evolved in the mammalian lineage as a way to regulate maternally and paternally expressed genes in the placenta during pregnancy and modulate metabolic functions related to growth, where the parental interests may be in conflict – paternal alleles favoring growth of the fetus at the expense of the mother while maternal alleles favor restricting resources to the fetus to ensure preservation of her nutritional needs^3,19,22^.Our data show some effects that are consistent with this theory. For example, three independent paternally inherited alleles on chromosome 1 are associated with increased LDL-C (**Fig. 2**) and total cholesterol (**Supplementary Fig. 7**); a paternal allele on chromosome 13 is also associated with increased systolic blood pressure (**Supplementary Fig. 6**). However, it is not always possible to interpret our results in light of this model, such as the association of maternal allele on chromosome 2 with decreased CIMT (**Supplementary Fig. 3**), or the maternal allele on chromosome 16 associated with decreased age of menarche (**Fig. 1**), which confers increased cardiovascular risk^23^. However, because many of the traits associated with POEs in this study were measured in adults, and none were measured in neonates, we are likely observing the downstream effects of processes that occurred *in utero*. Nonetheless, this kinship theory, or parent-conflict hypothesis, could account for the enrichment of parent of origin associations, particularly those with opposite effects, among metabolic and CVD-associated traits^1^.

Finally, we note that the parent of origin GWAS for 21 phenotypes in the Hutterites revealed overall twice as many genome-wide significant loci compared to standard GWAS of the same phenotypes in the same individuals, suggesting that variation at imprinted loci may represent some of the “missing heritability” of these phenotypes and potentially for the disease for which they confer risk. This idea is consistent with observations in mice showing that POEs contribute disproportionally to the heritability of 97 traits, including those related to total cholesterol, weight, HDL, and triglycerides^24^. Exactly how much loci with POEs in humans contribute to phenotypic variation and disease risk overall remains to be determined, but our study provides compelling evidence that it is likely to be significant for many important traits.

## SUBJECTS AND METHODS

### Sample Composition

The individuals in this study have participated in one or more of our studies on the genetics of complex traits in the Hutterites^25^^-^^27^. The more than 1,500 Hutterites in our study are related to each other in a 13-generation pedigree including 3,671 individuals.

### Genotype Data

Variants detected in the whole genome sequences of 98 Hutterites were previously imputed to an additional 1,317 individuals using PRIMAL, a high-accuracy pedigree based imputation method^28^. PRIMAL was used to phase alleles and assign parent of origin for 83% of about 12 million autosomal SNPs. For these studies, we selected SNPs that had a MAF >1% and genotype call rate > 85%. This yielded 5,891,982 autosomal SNPs. Parent of origin allele call rates differed among individuals and between phenotypes (**Supplementary Table 1**).

### Phenotype Data

We included 21 quantitative phenotypes that were previously measured in the Hutterites. Descriptions for each phenotype, as well as exclusion criteria, transformations, and covariates used with each phenotype in the GWAS, are available in the Supplementary Methods (**Supplementary Table 1**). Descriptions for 18 of the 21 phenotypes can be found in Cusanovich *et al.*^25^ The remaining three are described here. Height was measured in cm on a stadiometer with shoes removed. BMI was calculated using weight (kg, measured on scale) divided by height (m) squared. Age at menarche was collected retrospectively by interview.

### GWAS

We used a linear mixed model as implemented in GEMMA to test for genome wide association with 21 phenotypes using an additive model. We corrected for relatedness, as well as relevant covariates (**Supplementary Table 1**).

### Maternal and Paternal GWAS

To evaluated the evidence for POEs, we tested maternal and paternal alleles separately with each phenotype, comparing phenotypic differences between the maternally inherited alleles and between the paternally inherited alleles. We used a linear mixed model as implemented in GEMMA, which allows us to correct for relatedness as a random effect, as well as sex, age, and other covariates as fixed effects^29^. The linear mixed model for the parent of origin GWAS for testing maternal alleles and paternal alleles is shown in Equation 1 and Equation 2, respectively.
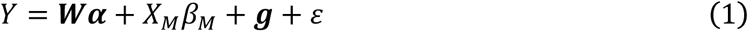

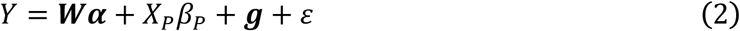

*n* is the number of individuals, *Y* is an *n* × 1 vector of quantitative traits, ***W*** is an *n* × *c* matrix of covariates (fixed effects) including intercept 1. ***α*** is a *c* × 1 vector of covariate coefficients. *X_M_* is an *n* × 1 vector of maternal alleles, and *X_P_* an *n* × 1 vector of paternal alleles. *β_M_* and *β_P_* are the effect sizes of maternal and paternal alleles, respectively. *g* is a vector of genetic effects with *g* ~ *N*(0, ***A****σ_g_*^2^) where ***A*** is the genetic relatedness matrix; *ε* is a vector of non-genetic effects with *ε* ~ *N*(0, ***I****σ_e_*^2^).

### Differential Effect GWAS (PO-GWAS)

To test for a difference in the same allele inherited from each parent, including opposite effects, we re-parameterized the test model (Equation 3) from Garg *et al.*^8^. The null model (Equation 4) is a standard GWAS model, ignoring parent of origin of alleles. The test model (Equation 3) is more significant when maternal and paternal alleles have differential effects on gene expression.
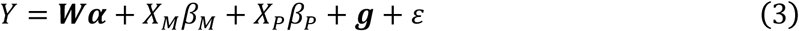

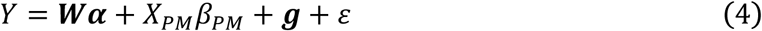

This new model allows us to measure the difference in parental effect of the same allele when the genotype is a covariate in Equation 5.
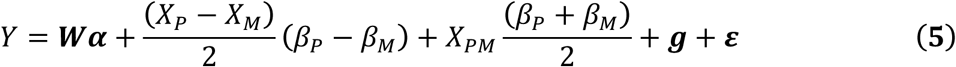

*X_PM_* is a *n* × 1 vector of genotypes with possible values [0, 1, 2], equivalent to *X_P_* + *X_M_*. (*β_P_* − *β_M_*) is the difference in parental effect size. If the difference in parental effect size is large and significantly different from 0 it suggests a parent of origin effect exists at this variant. 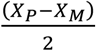 is a vector of genotypes with possible values [−1,0,1]. 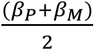 is the average parental effect size that is captured in normal GWAS using genotypes. The average genotypes are added in as a covariate, with the average parental effect size the corresponding covariate coefficient. This differential effect GWAS was tested in GEMMA using BIMBAM format to use average genotype values^30^.

### Parent of Origin eQTL Studies

RNA-seq data from LCLs were available from a previous study in the Hutterites^25^. For this study, sequencing reads were reprocessed as follows. Reads were trimmed for adaptors using Cutadapt (with reads <5 bp discarded) then remapped to hg19 using STAR indexed with gencode version 19 gene annotations^31,32^. To remove mapping bias, reads were processed using WASP mapping pipeline^33^. Gene counts were collected using HTSeq-count^34^. VerifyBamID was used to identify sample swaps to include individuals that were previously excluded^35^. Genes mapping to the X and Y chromosome were removed; genes with a Counts Per Million (CPM) value of 1 (expressed with less than 20 counts in the sample with lowest sequencing depth) were also removed. Limma was used to normalize and convert counts to log transformed CPM values ^36^. Technical covariates that showed a significant association with any of the top principal components were regressed out (RNA Integrity Number and RNA concentration).

### Maternal and Paternal Parent of Origin eQTL

LCL RNA-seq data was used to test the single parent model for the most significant SNP from the maternal or paternal only GWAS for each phenotype. We selected all genes detected as expressed in the LCLs and residing within 1Mb of each most significant associated SNP. Summary of the SNPs and genes tested are in **Supplementary Table 3**.

### Differential Parent of Origin eQTL

LCL RNA-seq data was used to test the opposite effect model for the most significant SNP in each region that was associated with a phenotype in the parent of origin opposite effects GWAS. We selected all genes detected as expressed in the LCLs and residing within 1Mb of each associated SNP. Summary of the SNPs and genes tested are in **Supplementary Table 4**.

## Supplemental Data description

**Supplementary Table 1**. Phenotypes and sample composition

**Supplementary Table 2**. GWAS results for all variants with p-value < 5×10^−08^

**Supplementary Table 3.** Candidate Genes for Parent of Origin eQTL

**Supplementary Table 4.** Candidate Genes for Parent of Origin Differential Effect eQTL

**Supplementary Table 5a-c**. Parent of Origin GWAS results with p-value < 5×10^−08^. Includes Maternal and Paternal GWAS and Differential Effect GWAS

**Supplementary Figure 1.** Manhattan and QQ Plots from Standard GWAS of 21 Quantitative Phenotypes

**Supplementary Figures 2-6**. Maternal and Paternal GWAS results for CIMT, LVMI, FEV1, LVMI, SBP, and Total Cholesterol

**Supplementary Figure 7**. Manhattan and QQ Plots from Differential Effect GWAS of 21 Quantitative Phenotypes

**Supplementary Figures 8-14**. Differential Parent of Origin GWAS results for LDL, LVMI, Triglycerides, Total Cholesterol, Blood Eosinophil Count, FEV1, and Neutrophil Count

**Supplementary Figure 15**. Differential Effect eQTL for rs7033776

## Acknowledgments

We thank Catherine Stanhope for help with processing phenotype data, Mark Abney and members of the Ober lab for useful discussions, Joe Urbanski and Lorenzo Pesce for assistance using Beagle, the many members of our field trip teams for help in phenotyping and collecting and processing samples, and the Hutterites for their continued support of our studies. This work was supported by NIH grants HL085197 and HD21244; and in part by NIH through resources provided by the Computation Institute and the Biological Sciences Division of the University of Chicago and Argonne National Laboratory, under grant 1S10OD018495-01. S.V.M was supported by NIH Grant T32 GM007197 and the Ruth L. Kirschstein NRSA Award F31HL134315.

## Web Resources

Code for PO-GWAS: https://github.com/smozaffari/POGWAS

## Author Contribution

S.V.M., D.L.N., and C.O. designed the study and wrote the paper. J.M.D., S.J.S., and R.M.L provided clinical data. S.V.M. performed analyses. All authors discussed results and commented on the manuscript.

## References

1. Peters, J. The role of genomic imprinting in biology and disease: an expanding view. Nature Reviews Genetics 15, 517–530 (2014).

2. Falls, J. G., Pulford, D. J., Wylie, A. A. & Jirtle, R. L. Genomic imprinting: implications for human disease. Am. J. Pathol. 154, 635–647 (1999).

3. Patten, M. M., Cowley, M., Oakey, R. J. & Feil, R. Regulatory links between imprinted genes: evolutionary predictions and consequences. Proc. Biol. Sci. 283, (2016).

4. Gabory, A. et al. H19 acts as a trans regulator of the imprinted gene network controlling growth in mice. Development 136, 3413–3421 (2009).

5. Varrault, A. et al. Zac1 regulates an imprinted gene network critically involved in the control of embryonic growth. Developmental Cell 11, 711–722 (2006).

6. Benonisdottir, S. et al. Epigenetic and genetic components of height regulation. Nat Comms 7, 13490 (2016).

7. Kong, A. et al. Parental origin of sequence variants associated with complex diseases. Nature Publishing Group 462, 868–874 (2009).

8. Garg, P., Borel, C. & Sharp, A. J. Detection of Parent-of-Origin Specific Expression Quantitative Trait Loci by Cis-Association Analysis of Gene Expression in Trios. PLoS ONE 7, e41695 (2012).

9. Perry, J. R. B. et al. Parent-of-origin-specific allelic associations among 106 genomic loci for age at menarche. Nature Publishing Group 514, 92–97 (2014).

10. Zoledziewska, M. et al. Height-reducing variants and selection for short stature in Sardinia. Nat Genet 47, 1352–1356 (2015).

11. Livne, O. E. et al. PRIMAL: Fast and Accurate Pedigree-based Imputation from Sequence Data in a Founder Population. PLoS Computational Biology 11, e1004139–14 (2015).

12. GTEx Consortium. Human genomics. The Genotype-Tissue Expression (GTEx) pilot analysis: multitissue gene regulation in humans. Science 348, 648–660 (2015).

13. Szafranski, P. et al. Pathogenetics of alveolar capillary dysplasia with misalignment of pulmonary veins. Human Genetics 135, 569–586 (2016).

14. Wang, Q., Lin, J. L.-C., Erives, A. J., Lin, C.-I. & Lin, J. J.-C. New insights into the roles of Xin repeat-containing proteins in cardiac development, function, and disease. Int Rev Cell Mol Biol 310, 89–128 (2014).

15. Nilsson, M. I. et al. Xin is a marker of skeletal muscle damage severity in myopathies. Am. J. Pathol. 183, 1703–1709 (2013).

16. O’Leary, N. A. et al. Reference sequence (RefSeq) database at NCBI: current status, taxonomic expansion, and functional annotation. Nucleic Acids Research 44, D733–45 (2016).

17. Brideau, C. M., Eilertson, K. E., Hagarman, J. A., Bustamante, C. D. & Soloway, P. D. Successful computational prediction of novel imprinted genes from epigenomic features. Mol. Cell. Biol. 30, 3357–3370 (2010).

18. Wang, Y. et al. From the Cover: Whole-genome association study identifies STK39 as a hypertension susceptibility gene. Proceedings of the National Academy of Sciences of the United States of America 106, 226–231 (2009).

19. Barlow, D. P. & Bartolomei, M. S. Genomic imprinting in mammals. Cold Spring Harb Perspect Biol 6, (2014).

20. Okae, H. et al. Re-investigation and RNA sequencing-based identification of genes with placenta-specific imprinted expression. Human Molecular Genetics 21, 548–558 (2012).

21. Proudhon, C. & Bourc’his, D. Identification and resolution of artifacts in the interpretation of imprinted gene expression. Brief Funct Genomics 9, 374–384 (2010).

22. Haig, D. The kinship theory of genomic imprinting. Annual review of ecology and systematics (2000).

23. Canoy, D. et al. Age at menarche and risks of coronary heart and other vascular diseases in a large UK cohort. Circulation 131, 237–244 (2015).

24. Mott, R. et al. The Architecture of Parent-of-Origin Effects in Mice. Cell 156, 332–342 (2014).

25. Cusanovich, D. A. et al. Integrated analyses of gene expression and genetic association studies in a founder population. Human Molecular Genetics 25, 2104–2112 (2016).

26. Weiss, L. A., Abney, M., Cook, E. H. & Ober, C. Sex-specific genetic architecture of whole blood serotonin levels. The American Journal of Human Genetics 76, 33–41(2005).

27. Abney, M., McPeek, M. S. & Ober, C. Broad and narrow heritabilities of quantitative traits in a founder population. The American Journal of Human Genetics 68, 1302–1307 (2001).

28. Livne, O. E. et al. PRIMAL: Fast and Accurate Pedigree-based Imputation from Sequence Data in a Founder Population. PLoS Computational Biology 11, e1004139–14 (2015).

29. Zhou, X. & Stephens, M. Genome-wide efficient mixed-model analysis for association studies. Nat Genet 44, 821–824 (2012).

30. Servin, B. & Stephens, M. Imputation-based analysis of association studies: candidate regions and quantitative traits. PLoS Genet 3, e114 (2007).

31. Dobin, A. et al. STAR: ultrafast universal RNA-seq aligner. Bioinformatics 29, 15–21 (2013).

32. Martin, M. Cutadapt removes adapter sequences from high-throughput sequencing reads. EMBnetjournal 17, pp. 10–12 (2011).

33. van de Geijn, B., McVicker, G., Gilad, Y. & Pritchard, J. K. WASP: allele-specific software for robust molecular quantitative trait locus discovery. Nat Meth 12, 1061–1063 (2015).

34. Anders, S., Pyl, P. T. & Huber, W. HTSeq–a Python framework to work with high-throughput sequencing data. Bioinformatics 31, 166–169 (2015).

35. Jun, G. et al. Detecting and estimating contamination of human DNA samples in sequencing and array-based genotype data. American journal of human genetics 91, 839–848 (2012).

36. Ritchie, M. E. et al. limma powers differential expression analyses for RNA-sequencing and microarray studies. Nucleic Acids Research 43, e47 (2015).

